# Assessing the predictive value of peak alpha frequency for the sensitivity to pain

**DOI:** 10.1101/2024.06.27.600974

**Authors:** Elisabeth S. May, Laura Tiemann, Cristina Gil Ávila, Felix S. Bott, Vanessa Hohn, Joachim Gross, Markus Ploner

## Abstract

Pain perception varies considerably between and within individuals. How the brain determines these variations has yet to be fully understood. The peak frequency of alpha oscillations (PAF) has recently been shown to predict an individual’s sensitivity to longer-lasting experimental and clinical pain. PAF is, thus, discussed as a potential biomarker and novel target for neuromodulatory treatments of pain. Here, we scrutinized the generalizability of the relation between PAF and pain. We applied brief painful laser stimuli to 159 healthy participants and related inter- and intra-individual variations of pain perception to PAF measured with electroencephalography. Comprehensive multiverse analyses across two sessions did not provide consistent evidence for a predictive role of PAF for brief experimental pain. This indicates that the relationship between PAF and pain does not generalize to all types of pain and calls for a systematic exploration of the relationship between PAF, pain perception, and other neuropsychiatric symptoms.

## Introduction

Pain perception varies both between and within individuals, i.e., from person to person and from moment to moment^1–3^. The neural mechanisms underlying such inter- and intra-individual variations of pain are not fully known. Understanding these mechanisms promises basic insights into the brain mechanisms of pain and could help to define new targets for pain treatment, e.g., through neuromodulation approaches^4–6^. Moreover, as variations in pain sensitivity can predict the development of chronic pain^7,8^ and responses to pain treatments^9–11^, identifying the associated neural mechanisms might help to develop clinically useful biomarkers of pain^12^.

In the brain, the perception of pain is shaped by neuronal oscillations at different frequencies^13–15^. Specifically, the peak frequency of alpha oscillations (peak alpha frequency, PAF) in sensorimotor regions has been related to inter-individual variations of pain^16,17^. PAF (illustrated in Fig. 1) is measured using spectral analysis of brain activity recorded with electro- or magnetoencephalography (EEG/MEG). It is commonly defined as the frequency with the highest peak in the alpha band (8-13 Hz)^18^. In previous studies, a slower resting-state PAF was associated with higher pain ratings during experimental pain lasting from minutes to weeks^19–21^, but see ^22^ for a positive relation between PAF and pain. Most recently, these findings have been extended to clinical conditions. In people with lung cancer, pre-operative resting-state PAF predicted pain severity after surgery^23^.

**Figure 1.**
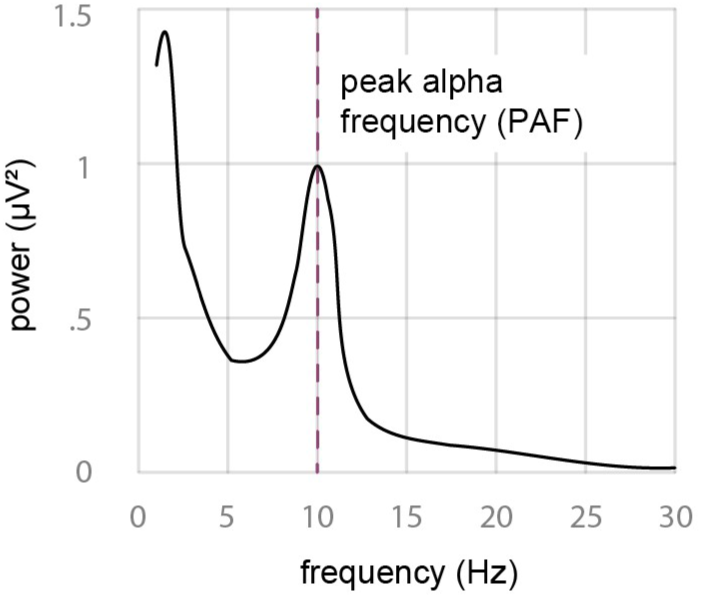
Peak alpha frequency (PAF). PAF is derived from the power spectrum of an individual’s electro- or magnetoencephalography (EEG/MEG) recording and is defined as the frequency with the highest power and a peak within the alpha range (8-13 Hz). As to the exact computation of PAF, different approaches exist.

**Figure 2.**
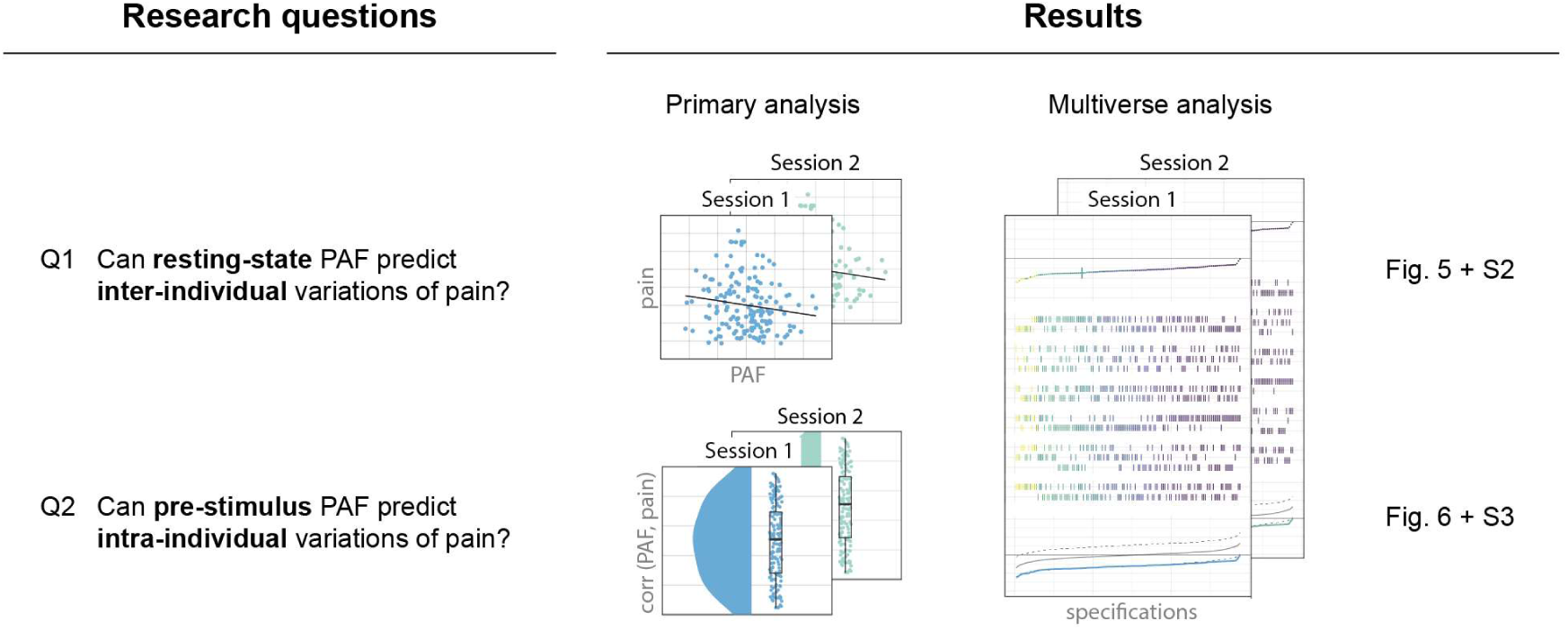
Outline of research questions, associated analyses, and results.

However, it is unclear how fundamental and generalizable the relationship between PAF and pain is. For instance, it is unknown whether the relationship similarly applies to different types and durations of experimental and chronic pain. Moreover, it is unknown whether PAF plays a similar role for both inter-individual, person-to-person variations of pain and intra-individual, moment-to-moment variations of pain. Understanding how generalizable the relationship between PAF and pain perception is, would help to assess the potential of PAF as a pain sensitivity biomarker. Moreover, such insights could have implications for other modalities and disorders. For instance, a faster PAF has been related to better cognitive performance^24–26^, and a slowed PAF has been observed in aging^26–30^, schizophrenia^31,32^, psychosis^33,34^, and dementia^35,36^. Thus, PAF might not only be a determinant of pain perception but a general determinant of perception, cognition, and susceptibility to neuropsychiatric disorders.

To better understand the role of PAF in the perception of pain, we related PAF to inter- and intra-individual variations in the perception of brief experimental pain stimuli in 159 healthy human participants. We specifically investigated whether resting-state PAF assessed by EEG can predict inter-individual variations of pain elicited by brief painful stimuli (Q1, Fig. 1). We further investigated whether prestimulus PAF can predict intra-individual variations of pain (Q2, Fig. 1). In a two-step approach, we performed primary analyses relying on an a priori chosen analytical pipeline followed by comprehensive multiverse analyses to test the robustness of results across different analytical settings^37,38^. Moreover, we investigated the replicability of findings across two recording sessions and confirmed the validity of the analytical approach in a positive control analysis.

## Results

One hundred fifty-nine healthy participants participated in two identical experimental sessions, separated by one month. The experimental paradigm is depicted in Figure 3. In each session, participants underwent a five-minute-long resting-state recording with eyes closed followed by an experimental pain paradigm. Eighty brief painful laser stimuli of 4 ms duration (DEKA Stimul 1340, Calenzano, Italy) were applied to the dorsum of the left hand at four different stimulus intensities. Participants verbally rated the perceived intensity of each stimulus on a scale ranging from 0 (“no pain”) to 100 (“maximally tolerable pain”). Brain activity was recorded using EEG with 64 active sensors placed according to the extended 10-20 system (Easycap, Wörthsee, Germany; Brain Products GmbH, Gilching, Germany) and automatically preprocessed^39,40^.

**Figure 3.**
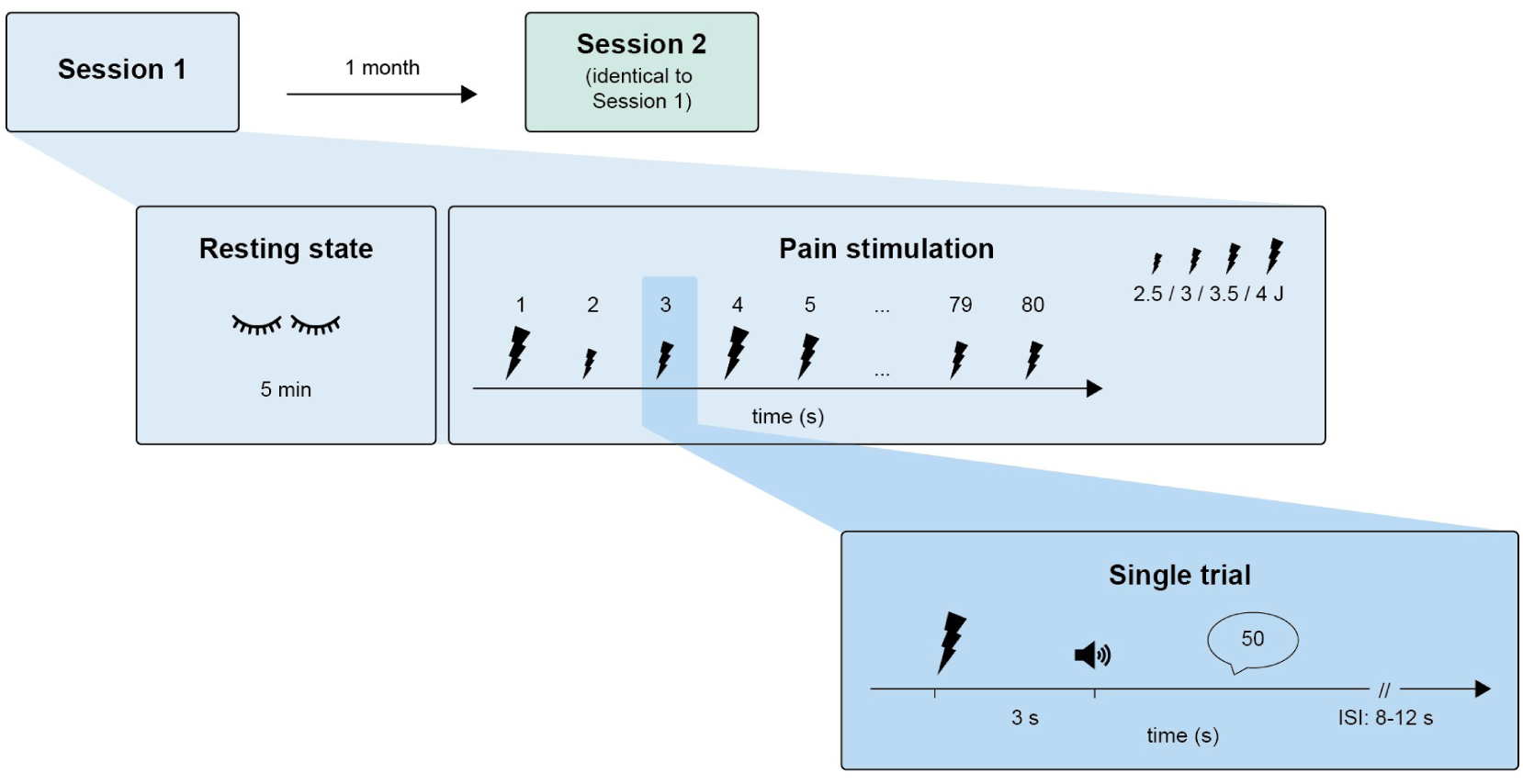
Paradigm. Participants took part in two identical experimental sessions separated by approximately one month. Each session started with a 5-minute eyes-closed resting-state recording. This was followed by a pain paradigm consisting of 80 laser stimuli with 4 ms duration and four fixed stimulus intensities (2.5, 3, 3.5, and 4 J) applied to the dorsum of the left hand in a pseudo-randomized order. Three seconds after each stimulus, an auditory cue prompted participants to verbally rate the perceived pain intensity on a scale from 0 (“no pain”) to 100 (“worst tolerable pain”). The inter-stimulus interval was 8 to 12 s. Brain activity was recorded using EEG. ISI, inter-stimulus interval.

PAF was estimated over the somatosensory cortex (S1), and inter- and intra-individual variations of S1 PAF were related to inter- and intra-individual variations of pain perception, respectively. As PAF can be estimated using various approaches^41,42^, we performed a multiverse analysis to investigate the robustness of our findings across different analytical approaches^37,43^. Thus, for each session, we performed a two-step approach. In a primary analysis, we first quantified S1 PAF and its relation to pain using an a priori chosen analytical pipeline. Subsequently, we complemented the primary analysis with a specification curve analysis, a type of multiverse analysis^37^. A specification thereby refers to a certain, single combination of analytical choices, i.e. one version of the analysis. Specification curve analysis performs multiple versions of the analysis, i.e., multiple specifications, and statistically tests the effects across all of them. All specifications performed here, including those of the primary analyses, are summarized in Table 1. To investigate the replicability of findings, we performed all analyses for sessions 1 and 2. The analysis was pre-registered (https://osf.io/r5buf).

**Table 1.**
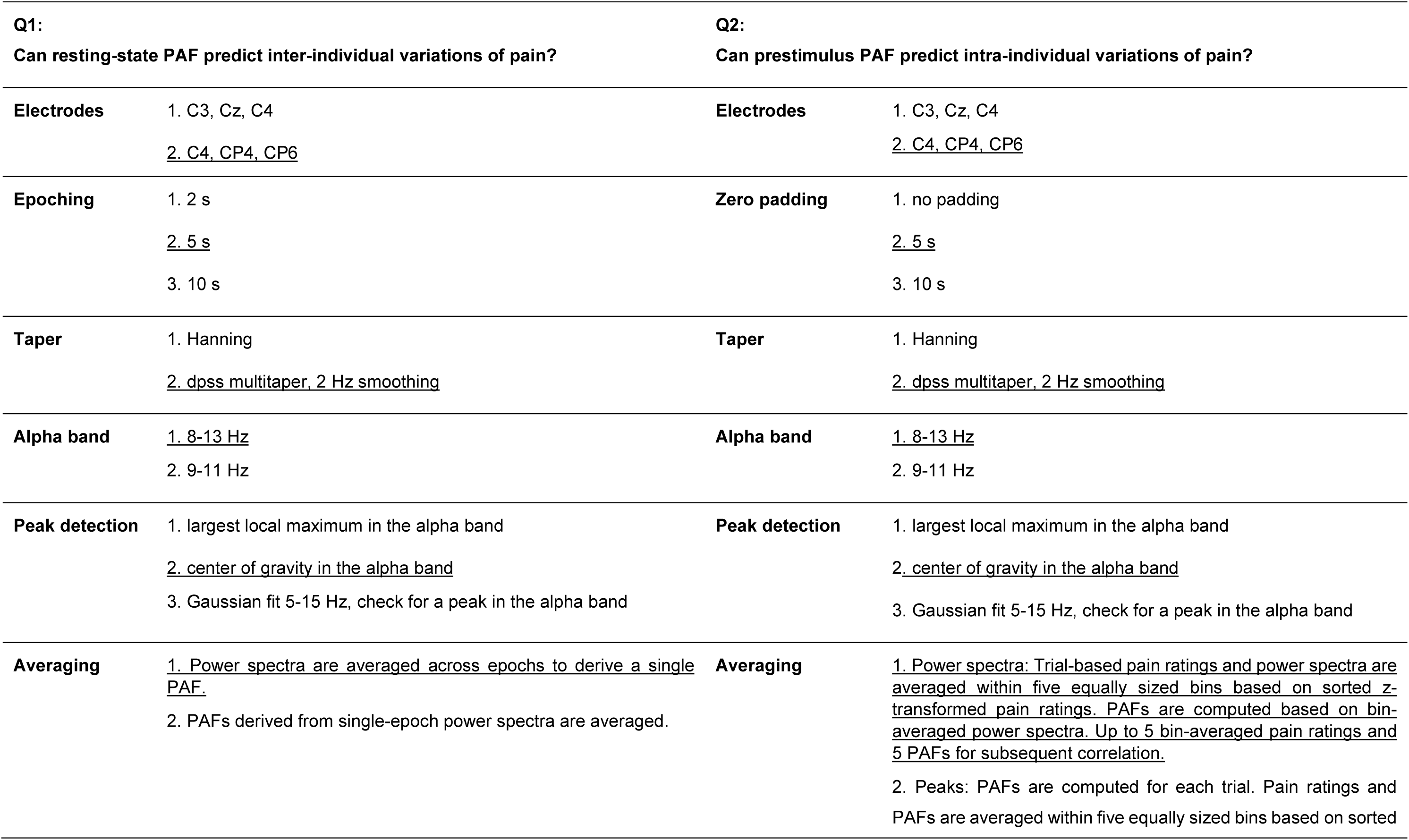
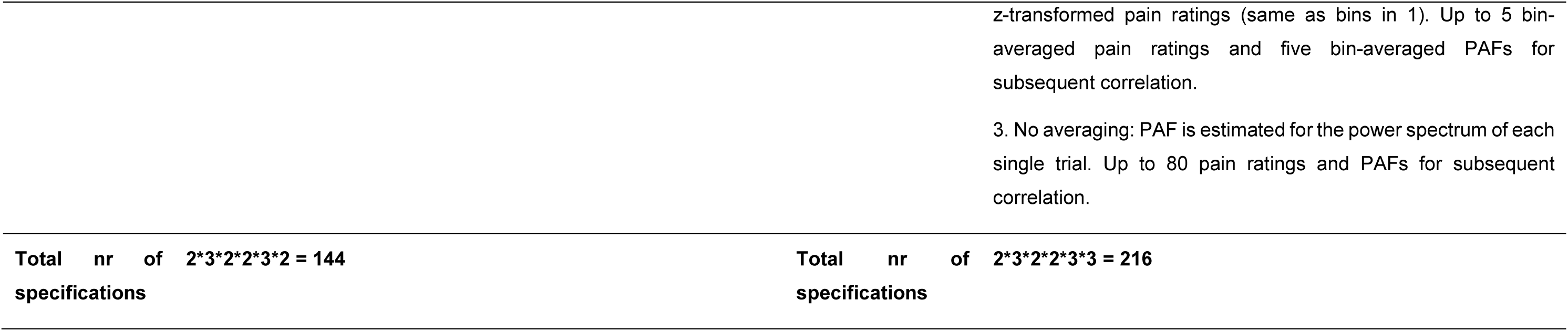
Analytical options for specification curve analyses. Settings used for primary analyses are underlined.

### Inter- and intra-individual variations of pain perception

We first confirmed that pain ratings in response to laser stimuli varied both between and within individuals (Fig. 4). The left panel of Figure 4A shows inter-individual variations of pain perception for both sessions. Across participants, the average pain rating across all trials was 22 ± 15 (mean ± standard deviation (SD), ranging from 0 to 62) for session 1 and 22 ± 14 (1 to 69) for session 2. The left panel of Figure 4B shows intra-individual variations of pain perception in terms of average SDs of pain ratings across the four stimulus intensities. The mean SD across participants was 11 ± 6 (0 to 30) for session 1 and 10 ± 6 (1 to 30) for session 2.

**Figure 4.**
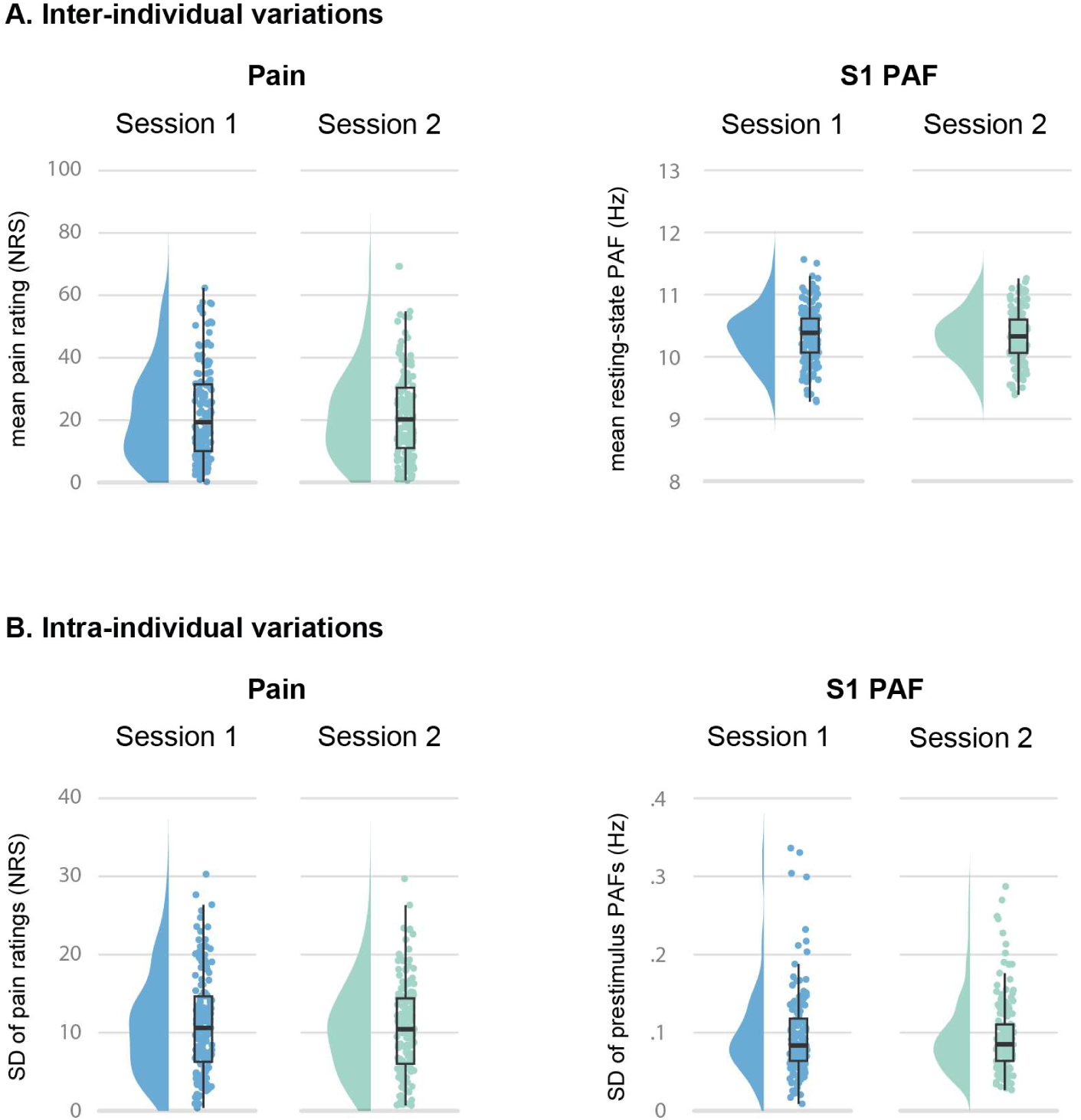
Inter- and intra-individual variations of pain perception and S1 PAF. A. Single-participant mean pain ratings and mean S1 resting-state PAFs obtained during primary analyses for sessions 1 and 2. To obtain mean pain ratings, ratings were first averaged separately for trials of the four different stimulus intensities and then averaged across them. Mean S1 PAFs were obtained by averaging PAF across 5 bins based on sorted pain ratings (see below). B. Single-participant SDs of pain ratings and S1 prestimulus PAFs obtained during primary analyses for sessions 1 and 2. SDs of pain ratings were calculated separately for trials of the four stimulus intensities and then averaged across them. SDs of S1 PAFs were calculated across bins. Raincloud plots show un-mirrored violin plots displaying the probability density function of the data, boxplots, and individual data points. Boxplots depict the sample median, first (Q1), and third quartiles (Q3). Whiskers extend from Q1 to the smallest value within Q1 - 1.5* interquartile range (IQR) and from Q3 to the largest values within Q3 + 1.5* IQR. SD, standard deviation; NRS, numerical rating scale.

Bayesian linear regressions revealed that mean pain ratings and SDs of pain ratings were highly correlated between the two sessions (Suppl. Fig. 1; mean pain ratings: R = .70, BF10 = 1.40 e+22; SDs of pain ratings: R = .78, BF10 = 2.66 e+30). Thus, pain ratings and their variability were stable across the two sessions one month apart.

### Inter- and intra-individual variations of S1 PAF

We next quantified how S1 PAF varied between and within individuals using PAF estimates from the primary analysis (Fig. 4, Table 1). Inter-individual variations of S1 PAF were measured using resting-state data. The mean S1 PAF (mean ± SD (range)) was 10.35 ± .44 Hz (9.28 to 11.26 Hz) for session 1 and 10.33 ± .40 Hz (9.39 to 11.57 Hz) for session 2 (Fig. 4A, right panel). Intra-individual variations of S1 PAF were derived from 2 s epochs preceding each stimulus during pain stimulation. The mean SD of S1 PAFs for the primary analysis was .10 ± .05 Hz (.01 to .34 Hz) for session 1 and .10 ± .05 Hz (.03 to .29 Hz) for session 2 (Fig. 4B, right panel). Thus, variations of S1 PAF within participants were substantially smaller than variations between participants^44,45^.

We next assessed the stability of S1 PAF across the two sessions. Mean S1 resting-state PAFs were strongly correlated between sessions, as shown by a Bayesian linear regression coefficient of R = .89 with a Bayes factor of 5 e+53 (Suppl. Fig. 1). The relation of SDs of S1 prestimulus PAFs between sessions was smaller, but still showed decisive evidence in favor of a relationship (Suppl. Fig. 1; R = .33, BF_10_ = 1109). Thus, S1 PAF and S1 PAF variability were stable across the two sessions one month apart.

### Relationship between PAF and inter-individual variations of pain perception

We further investigated whether S1 resting-state PAF can predict inter-individual variations of the perception of brief experimental pain. To this end, we related S1 resting-state PAF to average pain ratings across participants using Bayesian linear correlation coefficients, while controlling for the effects of age. This was done independently for both sessions using predefined settings for the primary analysis and 144 variations of the analysis for specification curve analysis (Table 1). For all specification curve analyses, results are first plotted in a so-called *descriptive* specification curve, which displays results and analytical settings of all specifications sorted according to the obtained effect size. Subsequently, based on *inferential* specification curve analyses^37^, we report and plot the results of three statistical tests that investigate the consistency and significance of findings across all specifications. These tests were based on (1) the median effect size across all specifications, (2) the share of significant specifications with the hypothesized negative correlation between S1 PAF and pain, and (3) the aggregated z-scores associated with the p-values of all specifications.

The primary analysis of session 1 yielded inconclusive evidence regarding the inter-individual relationship between S1 PAF and pain perception (Fig. 5A; R = -.15, BF_10_ = 1.04). In the multiverse analysis, the descriptive specification curve (Fig. 5B, top panel) showed negative relations between S1 PAF and pain for all 144 specifications. The strongest correlations were found for the central electrode selection (C3, Cz, C4) when determining S1 PAF on averaged power spectra (Fig. 5B, middle panel). In the inferential specification curve analysis (Fig. 5B, bottom panel), the observed specification curve was at the border of the one-sided 95 % confidence interval based on randomly permuted data. Accordingly, statistical tests across all specifications yielded p-values at the border of significance (*p median/share/aggr* = .054/.048/.04). These findings could not be replicated for session 2. For this session, the primary analysis showed strong evidence against an inter-individual relationship between S1 resting-state PAF and pain perception (Fig. 5A; R = .06, BF_10_ = .06). Moreover, the descriptive specification curve entailed about as many negative as positive correlations (Fig. 5B, top panel), and no statistical test of the inferential specification curve analysis was significant (Fig. 5B, bottom panel; *p median/share/aggr* = .56/.15/.56).

**Figure 5.**
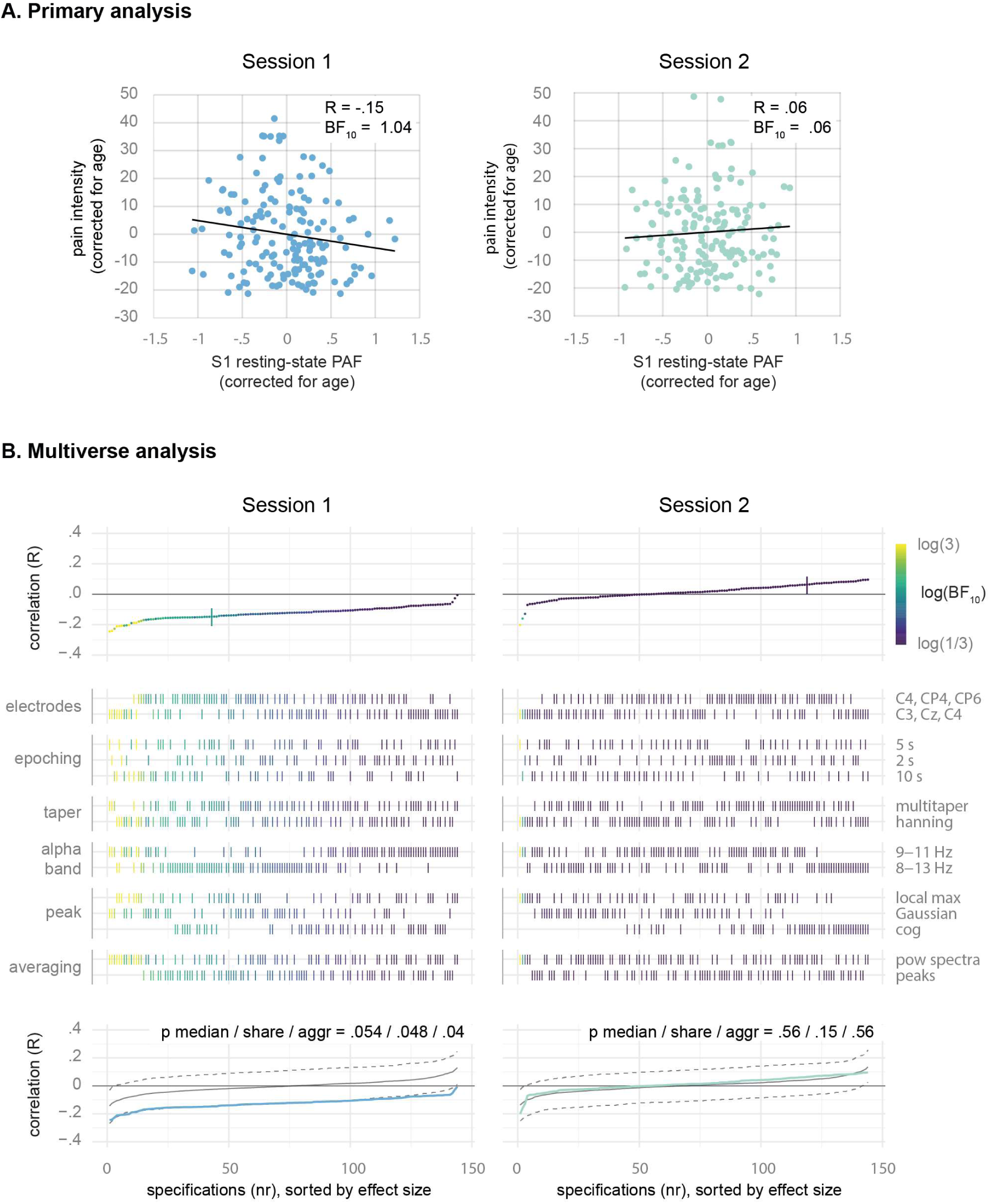
The relationship between S1 resting-state PAF and inter-individual variations of pain. A. Primary analysis. Scatter plots show the results of Bayesian linear correlations between S1 resting-state PAF and pain intensity, corrected for age. B. Multiverse analysis. The top panels show Pearson correlation coefficients obtained for all 144 specifications, sorted by effect size. The primary analysis is highlighted by a vertical bar. Middle panels mark the corresponding analytical choices of every specification (see Table 1). Results are color-coded based on Bayes factors. The scale’s upper and lower ends indicate moderate evidence for and against a relationship. The bottom panels display the inferential specification curve analysis showing the observed specification curve and the median and one-sided 95 % confidence intervals of 500 specification curves obtained from permutations of the data under the null hypothesis, all sorted by effect size. BF, Bayes factor; cog, center of gravity; PAF, peak alpha frequency; R, Pearson correlation coefficient; S1, primary somatosensory cortex.

To investigate effects beyond S1, we additionally examined the relation between global PAF and inter-individual variations of pain by averaging power spectra for PAF determination across all electrodes (Supp. Fig. 2). Aside from the electrode selection, the same analytical approach and settings were used, resulting in 72 specifications. For the global PAF, the primary analysis of session 1 provided inconclusive evidence regarding a relation to inter-individual variations of pain perception (Supp. Fig. 2A; R = -.12, BF_10_ = .56). The descriptive specification curve analysis revealed mostly negative correlations (Supp. Fig. 2B, top panel). In the inferential specification curve analysis, however, the observed curve was within the 95 % confidence interval, and all statistical tests showed negative results (Supp. Fig. 2B, bottom panel; *p median/share/aggr* = .10/1/.09). These findings were replicated in session 2 (primary analysis: R = .04, BF_10_ = .07; multiverse analysis: *p median/share/aggr* = .27/1/.23).

Together, these findings do not provide consistent evidence for a relationship between S1 PAF or global PAF and inter-individual variations of the perception of brief experimental pain stimuli.

### Relationship between PAF and intra-individual variations of pain perception

Next, we investigated whether prestimulus PAF could predict intra-individual variations of pain perception. This was based on single-trial pain ratings and PAFs estimated for 2 s prestimulus epochs immediately preceding the laser stimuli applied in the experimental pain paradigm. Intra-individual relationships between prestimulus PAF and pain ratings were quantified across trials using single-participant Pearson correlation coefficients. To assess whether these values differed from 0 across participants, we performed Bayesian t-tests. Again, relations were evaluated for both sessions using predefined settings for the primary analyses and 216 variations of analytical choices in specification curve analyses (Table 1).

Primary analyses of both sessions provided moderate evidence against a relation between S1 PAF and pain perception (Fig. 6A; session 1: BF_10_ = .12, session 2: BF_10_ = .19). In the multiverse analyses of both sessions, no specification provided evidence in favor of a relation between S1 prestimulus PAF and pain (Fig. 6B, top panels). Correspondingly, the observed curves in the inferential specification curve analyses were within the 95 % confidence intervals, and statistical tests resulted in insignificant p-values for both sessions (Fig. 6B, bottom panels; session 1: *p median/share/aggr* = .10/1/.076, session 2: *p median/share/aggr* = .21/1/.30).

**Figure 6.**
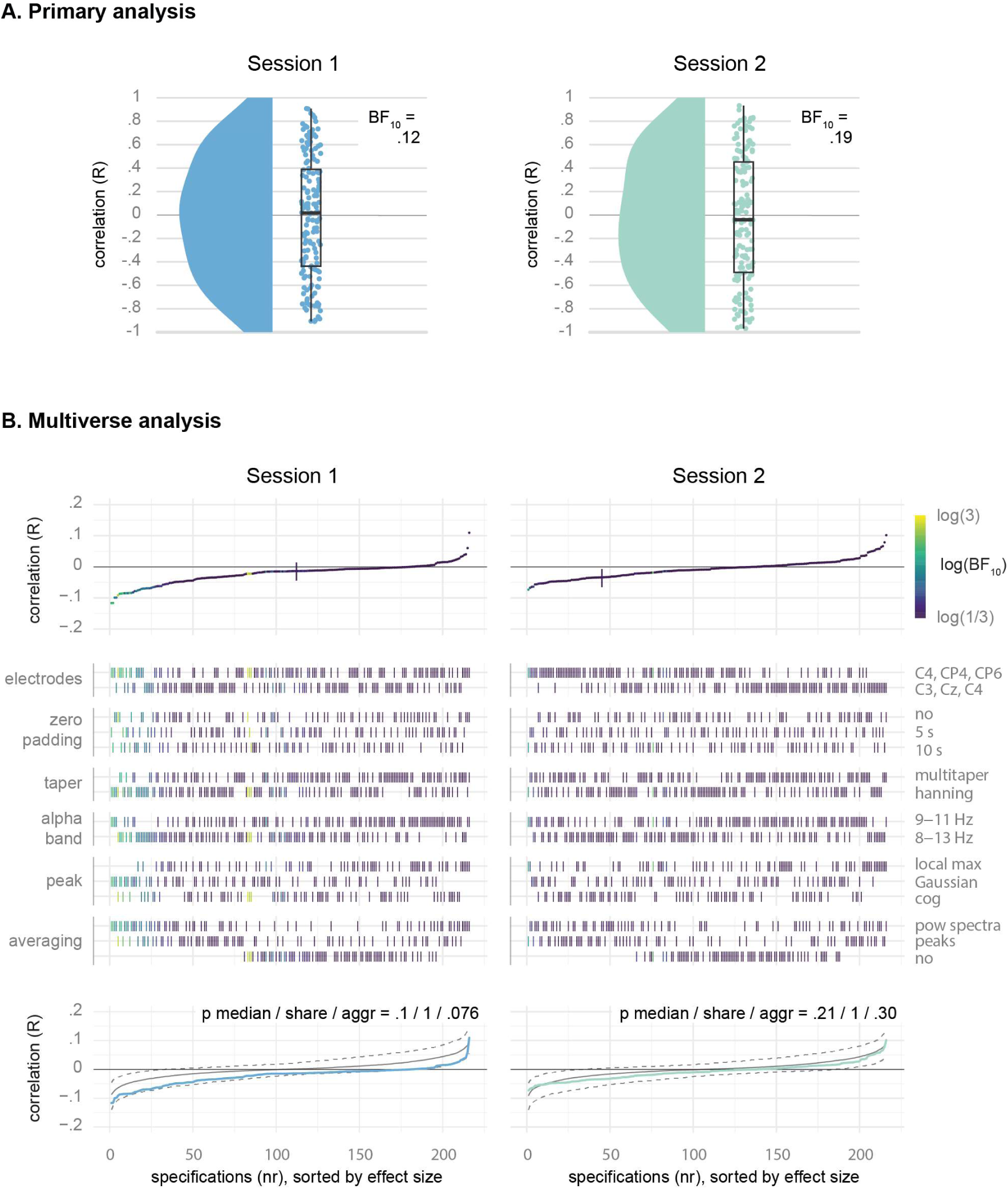
The relationship between S1 prestimulus PAF and intra-individual variations of pain. A. Primary analysis. Raincloud plots show single-participant correlation coefficients of the relation between S1 prestimulus PAF and pain intensity. Bayes factors are obtained from Bayesian t-tests of correlation coefficients vs. 0. B. Multiverse analysis. The top panels show Pearson correlation coefficients obtained for all 216 specifications, sorted by effect size. The primary analysis is highlighted by a vertical bar. Middle panels mark the corresponding analytical choices of every specification (see Table 1). Results are color-coded based on Bayes factors. The scale’s upper and lower ends indicate moderate evidence for and against a relationship. The bottom panels display the inferential specification curve analysis showing the observed specification curve and the median and one-sided 95 % confidence intervals of 500 specification curves obtained from permutations of the data under the null hypothesis, all sorted by effect size. BF, Bayes factor; cog, center of gravity; PAF, peak alpha frequency; R, Pearson correlation coefficient; S1, primary somatosensory cortex.

Beyond S1, results of primary analyses of global prestimulus PAF also showed strong evidence against a relationship with intra-individual variations of pain perception for both sessions (Supp. Fig. 3A; session 1: BF = .07, session 2: BF = .06), and none of 108 specifications showed evidence in favor of a relation (Supp. Fig. 3B, top panels). Finally, statistical tests across all specifications were consistently insignificant for both sessions (Supp. Fig. 3B, bottom panels; session 1: *p median/share/aggr* = .43/1/.37, session 2: *p median/share/aggr* =.70/1/.61).

Overall, these findings show replicable evidence against a relationship between prestimulus PAF and intra-individual variations of perceived pain.

### Relationship between PAF and age

As a positive control, we used our analytical approach to investigate the known inverse relation between resting-state PAF and age^26–30^. In these studies, PAF is usually estimated occipitally or globally. Thus, we repeated previous primary and multiverse analyses of the global resting-state PAF and related it to the age of participants.

Primary analyses (Supp. Fig. 4A) revealed inconclusive and moderate evidence against a relation between global PAF and age for session 1 (R = -.09, BF_10_ = .33) and session 2 (R = -.09, BF_10_ = .30). However, for both sessions, multiverse analyses with 72 specifications revealed specification curves outside of the 95 % confidence intervals and significant inverse relations between global resting-state PAF and age when quantifying effects across the entire specification curve (Supp. Fig. 4B; session 1: *p median/share/aggr* = .008/.004/.02, session 2: *p median/share/aggr* =.03/.02/.06). Thus, this positive control analysis shows that the current analytical approach is sensitive to previously reported findings by detecting a slowing of the global resting-state PAF with age across specifications.

## Discussion

The current study investigated the relationship between PAF and variations in the perception of brief experimental pain in a large sample of healthy human participants. We first asked whether resting-state PAF could predict inter-individual variations of pain. This question has been recently investigated using somatosensory PAF as a marker of sensitivity to pain lasting from minutes to weeks^19–21,23^. Here, we investigated it for brief experimental pain stimuli lasting milliseconds. Earlier findings of an inverse relation between PAF and pain perception could not be consistently replicated. Next, we investigated whether PAF immediately before a painful stimulus could predict intra-individual variations of pain perception elicited by that stimulus. Results from both sessions uniformly provided evidence against such a relation. Thus, based on the current results, PAF is not a general determinant of all types of pain.

Evidence for a relation between S1 PAF and inter-individual, person-to-person variations of pain perception was only found in session 1. Of the 144 performed specifications, nine provided evidence supporting this relation. However, this finding could not be replicated in session 2. Notably, there were no discernible differences between the two sessions. Experimental settings were identical, and mean pain ratings, S1 PAFs, and their variability were strongly correlated between sessions (Suppl. Fig. 1). Thus, the most feasible explanation for the disparity between the two sessions is that the relationship between S1 PAF and a person’s sensitivity to brief experimental pain is weak and variable. This emphasizes the importance of internal (like in the current study) and external replications, particularly in the context of potential clinical applications.

Concerning intra-individual, moment-to-moment variations of pain, the current results provide consistent evidence against a predictive role of S1 or global prestimulus PAF. Similar to previous studies^44,45^, intra-individual variability of PAF was substantially smaller than inter-individual variability. Furthermore, PAF was highly correlated between sessions (Suppl. Fig. 1), in line with a high within-participant stability of PAF over time^45–47^. One could deduce that intra-individual changes of PAF might be too small to be functionally relevant. However, previous studies did report task-related intra-individual changes of PAF, for example, in a working memory task^44^ or during hand immersion in painful hot water^48^.

The present study did not show an association between S1 PAF and inter- or intra-individual variations of pain elicited by brief noxious stimuli. This contrasts with previous studies showing that S1 PAF predicted inter-individual variations of pain perception lasting from minutes to weeks^19–21,23^. This disparity might indicate that PAF plays a stronger role in longer-lasting than brief pain. Intriguingly, the longer pain lasts, the more it is influenced by contextual factors like expectations and previous learning experiences^49–51^. In contrast, shorter pain is more closely related to objective sensory information and its cerebral processing^51^. Thus, the disparity between the present and previous findings might indicate that PAF might be more closely related to contextual than to sensory processes in the processing of pain. This would be in line with a role of PAF for more general cognitive processes and a higher-level susceptibility to pain and other neuropsychiatric conditions. Importantly, this lack of specificity does not limit the potential value of PAF as a clinically useful biomarker of pain sensitivity or as a novel target for pain treatment by using neuromodulatory approaches, such as repetitive transcranial magnetic stimulation^52^.

A potential limitation of the current work concerns the selection of analytical settings to estimate PAF. We did test the robustness of our findings across many analytical choices. However, in light of the multitude of analytical options used in the field, the selected specifications could not be completely exhaustive and, thus, were, to some extent, subjective^37^. However, we were able to detect the often-described slowing of global PAF with age using these specifications, confirming the sensitivity of our analytical approach. In addition, the current study focused on an extensive investigation of a single type of experimental pain in a large sample, including an internal replication. Future studies might systematically investigate how different types of experimental and clinical pain depend on PAF.

In conclusion, the current study demonstrates that the relation between PAF and pain is not generalizable across all types and durations of pain. Together with previous findings, the study highlights how the brain mechanisms of different types and durations of pain differ. Importantly, the findings do not argue against the predictive role of PAF as a marker for pain sensitivity and a potential target of neuromodulatory treatment approaches for chronic pain. However, they call for a systematic exploration of the relationship between PAF and the sensitivity to different types and durations of pain. Such a systematic exploration would help to better understand the perspectives and limits of PAF as a biomarker of pain and other neuropsychiatric conditions.

## Methods

### Dataset

The study uses a dataset recorded by the PainLabMunich (https://www.painlabmunich.de/) at the Technical University of Munich (TUM) in Germany between December 2019 and December 2022 (https://clinicaltrials.gov/ct2/show/NCT05616091). The study protocol was approved by the Ethics Committee of the Medical Faculty of the TUM and conducted following the latest version of the Declaration of Helsinki.

The present analysis was preregistered on the OSF REGISTRIES (https://osf.io/r5buf). At the time of preregistration, no previous analysis of the dataset had been performed, published, or presented. Thus, the authors had no prior knowledge about the dataset.

### Participants

We used non-probability sampling to recruit equal numbers of females and males and equal numbers of participants with ages <= 40 years and > 40 years. Participants were recruited using advertisements posted across the TUM and its university hospital and on corresponding web pages. In addition, participants from previous studies who had given their permission were contacted again. Following this approach, the sample included participants of a wide age range but primarily white men and women pursuing or previously having acquired higher education. Thus, the study sample constitutes a specific sample that is not necessarily representative of Germany or any region.

The present data analysis includes all participants for which resting-state and experimental pain recordings were available for both sessions. Of the total sample of n = 166, four participants did not return to the second session. For three other participants, one of the two resting-state recordings was not obtained due to technical errors. These participants were excluded from all analyses. Thus, the final sample size is n = 159 (80/79 female/male; age = 40 years ± 18 years (mean ± SD), range 18 to 86 years). Inclusion criteria were age ≥ 18 years and right-handedness. Exclusion criteria were pregnancy, neurological or psychiatric diseases (e.g. epilepsy, stroke, depression, anxiety disorders), severe general illnesses (e.g. tumors, diabetes), skin diseases (e.g. dermatitis, psoriasis, or eczema), current or recurrent pain, (regular) intake of centrally acting, antibiotic, or analgesic medication, previous surgical procedures involving the brain or spinal cord, past head trauma followed by impairment of consciousness, past fainting spells or syncopes, and side-effects following previous electrical, magnetic, or thermal stimulation.

### Procedure and paradigm

The paradigm is depicted in Figure 3. Participants took part in two identical experimental sessions on two separate days, scheduled about one month apart from each other. For female participants before menopause, recordings were always performed between days 5 and 10 of the menstrual cycle. For 153 participants, the inter-session interval was 29 ± 9 days (mean ± SD), ranging from 17 to 98 days. For the remaining 6 participants, the mean inter-session intervals were longer than 200 days (261 ± 34 days (mean ± SD), ranging from 223 to 308 days) due to recording restrictions during the COVID-19 pandemic.

In each session, healthy participants first completed clinical and demographic questionnaires (including questionnaires about sleep quality, anxiety and depression, positive and negative affect, and pain sensitivity). This was followed by familiarization with the laser stimulation and the pain rating procedure. To this end, 20 laser stimuli were applied in a pseudo-randomized sequence, with five stimuli at each of the four stimulation intensities which would be applied later during the experimental paradigm (see below). Subsequently, EEG caps were prepared. Participants then underwent a five-minute resting-state recording in which they were asked to stay relaxed but wakeful with their eyes closed. During the immediately following experimental pain paradigm, participants received 80 brief experimental pain stimuli at four different intensities to the dorsum of the left hand, again with their eyes closed. Prompted by an auditory cue 3 s after each stimulus, participants verbally rated the perceived intensity of each stimulus on a scale ranging from 0 (“no pain”) to 100 (“maximally tolerable pain”). During recordings, participants wore protective goggles and sat in a comfortable chair. White noise was played on headphones to eliminate the effects of any ambient noise.

Please see the preregistration of the initial study (https://clinicaltrials.gov/ct2/show/NCT05616091) for a more detailed description of the data collection procedure, including the collection of demographic and psychological variables and skin conductance responses, which are not analyzed here.

### Stimulation

Pain stimuli to the left hand were applied using a laser device (DEKA Stimul 1340, Calenzano, Italy) with a wavelength of 1340 nm, a duration of 4 ms, and a spot diameter of 7 mm. In a pseudo-randomized order, stimuli were applied at four fixed intensities (20 each at 2.5, 3, 3.5, and 4 J) with an inter-stimulus interval between 8 and 12 s. Unbeknown to the participant, stimuli were applied in 4 blocks of 20 stimuli, with 5 stimuli at each intensity. Within these blocks, the order was pseudo-randomized with the restriction that no more than two stimuli of the same intensity were applied in a row. The stimulation site was slightly changed after each stimulus to avoid tissue damage, habituation, and sensitization. After 40 trials, a short break was scheduled, during which the participants could open their eyes and adjust their seating position.

### Recordings and preprocessing

Brain activity was recorded using 64 actiCAP slim/snap sensors placed according to the extended international 10-20 system (Easycap, Wörthsee, Germany) and BrainAmp MR plus amplifiers (Brain Products GmbH, Gilching, Germany). During recordings, sensors were referenced to FCz and grounded at Fpz. Signals were sampled with a sampling frequency of 1000 Hz and band-pass-filtered between 0.016 and 250 Hz. Impedances were kept below 20 kΩ.

EEG data were automatically preprocessed separately for resting-state and pain stimulation recordings and sessions 1 and 2 using the automatic preprocessing part of the DISCOVER-EEG pipeline^39,40^ based on the MATLAB (Mathworks, Natick, MA) toolbox EEGLAB^53^. This included line noise removal, bad channel detection and interpolation, re-referencing to the average reference, independent component analysis (ICA), and the automatic detection of bad segments. For the two resting-state recordings, the continuous 5-minute recordings were preprocessed per session and later segmented into epochs of 2, 5, and 10 s length with 50 % overlap (see below). Participants were included in further analyses if at least five clean epochs were available after removal of epochs containing segments marked as bad during preprocessing (n = 159 for 2 and 5 s epoch lengths for sessions 1 and 2, n = 158 and 157 for 10 s epoch lengths of sessions 1 and 2, respectively). Numbers (mean ± SD) of remaining clean trials were 269 ± 41, 101 ± 20, and 45 ± 13 for 2 s, 5 s, and 10 s epochs in session 1 and 267 ± 40, 99 ± 21, and 45 ± 13 for 2 s, 5 s, and 10 s epochs in session 2, respectively.

For each of the two pain stimulation recordings, all 2 s prestimulus intervals immediately preceding the laser stimuli were appended before performing ICA and bad segment detection. Participants were included in further analyses if clean prestimulus intervals of at least five trials at each stimulus intensity in both sessions remained after removal of trials containing bad segments marked during preprocessing (n = 157). The mean numbers ± standard deviation (SD) of the remaining clean trials were 18 ± 3, 18 ± 3, 18 ± 3, and 18 ± 3 for stimulus intensities 1 to 4 of session 1, and 19 ± 1, 19 ± 2, 19 ± 2, and 19 ± 2 for stimulus intensities 1 to 4 of session 2, respectively.

### Analyses

We performed a two-step approach. First, we conducted a primary analysis by determining the PAF for a region of interest covering the primary somatosensory cortex (S1 PAF) using an a priori chosen analytical pipeline. Second, we complemented the primary analysis with a multiverse analysis, particularly specification curve analysis, which quantifies and statistically tests outcomes across a wide range of analytical options to investigate the robustness of findings^37^. For research questions 1 and 2 (Fig. 2), variations of S1 resting-state and prestimulus PAF were related to inter- and intraindvidiual variations of pain ratings following laser stimulation, respectively. All analyses were performed separately for sessions 1 and 2 to investigate the replicability of findings.

Analyses were performed using MATLAB (Mathworks, Natick, MA) with customized code and the MATLAB-based toolbox FieldTrip^54^. In addition, the R programming environment^55^, including the “BayesFactor”^56^ and “specr”^57^ packages, were used.

#### Question 1: Can resting-state PAF predict inter-individual variations of pain?

For the analysis of research question 1, single-participant S1 PAF estimates based on eyes-closed resting-state EEG recordings were related to single-participant mean pain ratings obtained during pain stimulation. Pain ratings were first averaged across trials at each stimulus intensity and then across the four intensities. Hence, one overall measure of pain perception was obtained per participant. To estimate S1 PAF, resting-state EEG data were segmented into epochs with 50 % overlap, and one single estimate of the S1 resting-state PAF was obtained for each participant. We then related average pain ratings to S1 PAF estimates across individuals using Bayesian linear correlations, while controlling for age (see below).

For the primary analysis, to obtain S1 PAF estimates, preprocessed data were segmented into 5 s epochs (for a 0.2 Hz spectral resolution) with 50 % overlap. Epochs containing segments marked as bad during preprocessing were rejected. For each epoch and channel, power spectra between 5 and 15 Hz were calculated using Fast Fourier Transformation and dpss multitapering with 2 Hz spectral smoothing. Estimates of S1 PAF were obtained by averaging power spectra across somatosensory channels of interest contralateral to stimulation (C4, CP4, and CP6). Then, power spectra were averaged across all epochs. Using these averaged power spectra, S1 PAF was determined per participant by calculating the center of gravity for an alpha band of interest between 8 and 13 Hz. The analytical settings chosen as primary analysis were based on previous literature.

For the multiverse analysis, several analytical settings were varied and results across different specifications of the analysis were analyzed using specification curve analysis^37^. One specification hereby refers to one combination of analytical settings, i.e. one version of the analysis. The chosen analytical variants were based on previous studies quantifying the PAF (including, amongst others, ^19–21,41,44,58^) as well as internal discussions and theoretical considerations of reasonable analytical choices. Table 1 presents an overview of all specifications selected for the current study, which included variations of selected electrodes of interest, epoch length, taper applied during power spectra calculations, alpha band of interest, PAF detection method, and the way of averaging power spectra vs. obtained PAFs per participant. The combination of all possible analytical decisions resulted in 144 specifications (analysis versions) for relating S1 PAF to inter-individual variations of pain pain, including the approach chosen as the primary analysis.

To investigate a potential relation between resting-state PAF and inter-individual variations of pain beyond S1, we additionally computed PAF on a global level based on power spectra averaged across all electrodes (global PAF). Aside from the electrode selection, identical primary and multiverse analyses to the ones implemented for S1 PAF were performed (Suppl. Fig. 2). This resulted in half of the specifications, i.e., 72 specifications, for the specification curve analysis of each session, since the electrode selection was not varied for global PAF.

At the stage of preregistering this analysis, we additionally planned to assess the stability of resting-state PAF as a marker of a participant’s pain sensitivity by using resting-state PAFs obtained from session 1 to predict average pain ratings in session 2 and vice versa (https://osf.io/r5buf). In light of our findings, which did not show a clear predictive value of S1 or global PAF for inter-individual pain, we omitted this analysis.

#### Question 2: Can prestimulus PAF predict intra-individual variations of pain?

Analyses for question 2 used data obtained during the experimental pain paradigm only and related single-trial S1 PAF estimates to single-trial pain ratings for both sessions. To minimize the influence of different stimulus intensities, the pain ratings of each participant were z-transformed per stimulus intensity before entering subsequent analyses^59^. S1 PAF estimates were derived from preprocessed 2 s prestimulus time windows immediately before each laser stimulus. Intra-individual relationships between S1 PAF and pain ratings were quantified using single-participant Pearson correlation coefficients, which then entered statistical analyses (see below).

For the primary analysis, 2 s prestimulus intervals were zero-padded to 5 s (obtaining a 0.2 Hz spectral resolution). In line with the primary analysis for question 1, power spectra between 5 and 15 Hz were calculated for each trial and channel using a Fast Fourier Transformation and dpss multitapering with 2 Hz spectral smoothing. Then, power spectra were averaged across channels C4, CP4, CP6. Subsequently, to increase the signal-to-noise ratio, trials were assigned to five bins based on associated pain ratings. Each bin comprised the same number of trails. The first bin included trials with pain ratings lower than the 20th percentile of all z-transformed pain ratings. The second bin included trials with pain ratings between the 20th and 40th percentile. This pattern continued for the other bins, each covering 20 % of the pain rating distribution. Both pain ratings and power spectra were then averaged within the five bins. Subsequently, S1 PAF was determined for each bin-averaged power spectrum by calculating the center of gravity between 8 and 13 Hz. Lastly, for each participant, the intra-individual relation between bin-based S1 PAFs and bin-averaged pain ratings was quantified with the Pearson correlation coefficient.

As before, specification curve analyses were performed to investigate the effects of different analytical options on relations between S1 PAF and pain^37^. These were identical to the variations performed for Q1 with modifications for zero-padding and averaging of S1 PAFs and pain ratings (Table 1). Combining all options resulted in 216 specifications for the intra-indvidual prediction of pain ratings by S1 PAF. Participants were only included in further analyses if the PAF of at least three bins / single trials (depending on the specification) could be determined for subsequent correlation (since there might not always be a local maximum or a peak of a Gaussian fit within the alpha band of interest).

As for Q1, effects beyond S1 were investigated by performing corresponding primary and specification curve analyses of the relation of global PAF to intra-individual variations of pain for both sessions (Suppl. Fig. 3). Power spectra were now averaged across all electrodes instead of somatosensory channels of interest, which resulted in 108 specifications for the prediction of pain ratings by global prestimulus PAF.

### Statistical analyses

#### Statistical power

To obtain an estimate of statistical power for our primary analyses, we performed Bayes factor (BF) design analyses (BFDA)^60^ using the R package “BFDA”^61^ for n = 159 participants with default priors and 10.000 simulations. To address the statistical power of question 1, a BFDA for a Bayesian linear correlation was performed. For a negative correlation of medium size R = -0.3 and BF_10_ boundaries of 1/3 and 3, BFDA revealed a statistical power of 93.6 % with a false positive rate of 0.9 %. Using more conservative BF_10_ boundaries of 1/10 and 10, statistical power was 85.8 % with 0.2 % false positives. To estimate statistical power for question 2, a BFDA was calculated for a Bayesian one-sided one-sample t-test. For a medium effect size of Cohen’s d = 0.5 and BF_10_ boundaries of 1/3 and 3, BFDA revealed a statistical power of 100 % with a false positive rate of 0.8 %. For more conservative BF_10_ boundaries of 1/10 and 10, statistical power remained at 100 % with 0.2 % false positives. Thus, the study was well-powered to detect medium effect sizes.

#### Statistical models

Our first hypothesis, associated with question 1, was that variations of an individual’s S1 resting-state PAF are inversely related to inter-individual variations of pain perception. For each specification, including the specification corresponding to the primary analysis, Bayesian linear correlation coefficients were calculated across participants using individual resting-state PAFs and average pain ratings. To control for the effects of age on PAF and pain, we employed a partial correlation approach by computing these Bayesian linear correlations between the residuals of two linear regression models predicting S1 PAF and pain from age, respectively.

For primary analyses, statistical inference is based on Bayes factors (BF_10_) obtained from Bayesian linear correlations. A BF_10_ > 3, > 10, and >100 (<1/3, < 1/10, < 100) indicates at least moderate, strong, and decisive evidence for (against) H1, respectively. For multiverse analyses, results across all specifications are summarized using descriptive specification curves^37^. To assess statistical significance across all specifications, inferential specification curve analyses were calculated, plotted, and complemented by three statistical tests based on a permutation approach as proposed by ^37^. Of note, these tests are based on frequentist statistical inference, since, to the best of our knowledge, Bayesian approaches to specification curve analysis do not exist yet. To this end, the individual average pain ratings were randomly permuted, and the analyses for all specifications repeated. This was done 500 times, followed by three statistical tests. In each test, a p-value was derived by comparing a test statistic of the original specification curve to the corresponding distribution of test statistics of randomized specification curves. The considered test statistics were 1.) the median effect size (i.e. correlation coefficient) across all specifications (*p median*), 2.) the share of specifications with the hypothesized negative correlation between resting-state PAF and pain and a BF_10_ > 3 (*p share*), and 3.) z-values associated with p-values summarized across all specifications according to Stouffer’s method (*p aggr*). The significance of the three tests was assessed with a one-sided p < 0.05. Note that to obtain the single-specification p-values for the third test, a frequentist Pearson correlation was computed for each specification, as before controlling for effects of age.

Question 2 was analyzed accordingly to address the second hypothesis that intra-individual variations of the PAF are inversely related to intra-individual variations of pain perception. Here, single-participant Pearson correlation coefficients obtained from all specifications including those from the primary analysis, were statistically tested using one-sided Bayesian t-tests, comparing the mean across correlation coefficients to 0.

As before, results from Bayesian t-tests of the primary analysis are interpreted based on corresponding Bayes factors. Subsequently, descriptive and inferential specification curve analyses were calculated in line with the statistical approach for question 1. Again, statistical significance across all specifications was assessed using the three frequentist tests outlined above based on the median effect size, the share of specifications with an effect in the predicted direction and a BF > 3, and the average z-value across all specifications. Here, for each of the 500 permutations, the assignment of single trial pain ratings and EEG data was shuffled for every participant.

### Control analysis

As a positive control analysis, we used our analytical approach to investigate the well-known slowing of the (mostly occipitally or globally) measured PAF throughout adulthood (e.g., ^26–30^). To this end, primary and multiverse analyses of the global resting-state PAF were repeated for both sessions with slight modifications (Suppl. Fig. 4). Rather than relating global resting-state PAF to average pain ratings using a partial correlation approach, it was now directly related to the age of participants using Bayesian linear correlations. This was done for all 72 specifications investigated in previous analyses of the global resting-state PAF. Statistical analyses and inferences remained the same, now shuffling the assignment of obtained PAF estimates and age across participants for the three statistical tests of the inferential specification curve analysis.

## Data availability

Raw and preprocessed data in standardized EEG-BIDS format^62^ are shared on osf.io (https://osf.io/s75q6/).

## Code availability

Code used for the current manuscript is openly available on osf.io (https://osf.io/s75q6/).

## Acknowledgements

We thank our student research assistants for their great help with data acquisition. The study has been supported by the TUM Innovation Network Neurotechnology for Mental Health (NEUROTECH) and the Deutsche Forschungsgemeinschaft (PL321/14-1, SFB1158).

## Author contributions

Conceptualization: ESM, LT, MP; Methodology: ESM, JG, MP; Software: ESM, CGA, FSB; Validation: ESM, CGA; Formal analysis: ESM; Investigation: LT; Resources: MP; Data curation: ESM, LT; Writing - original draft: ESM, MP; Writing – Review & Editing: ESM, LT, CGA, FSB, VH, JG, MP; Visualization: ESM, CGA, MP; Supervision: MP; Project administration: MP; Funding acquisition: MP.

## Competing interests

The authors declare no competing interests.

## Supplementary material

**Supplementary Figure 1.**
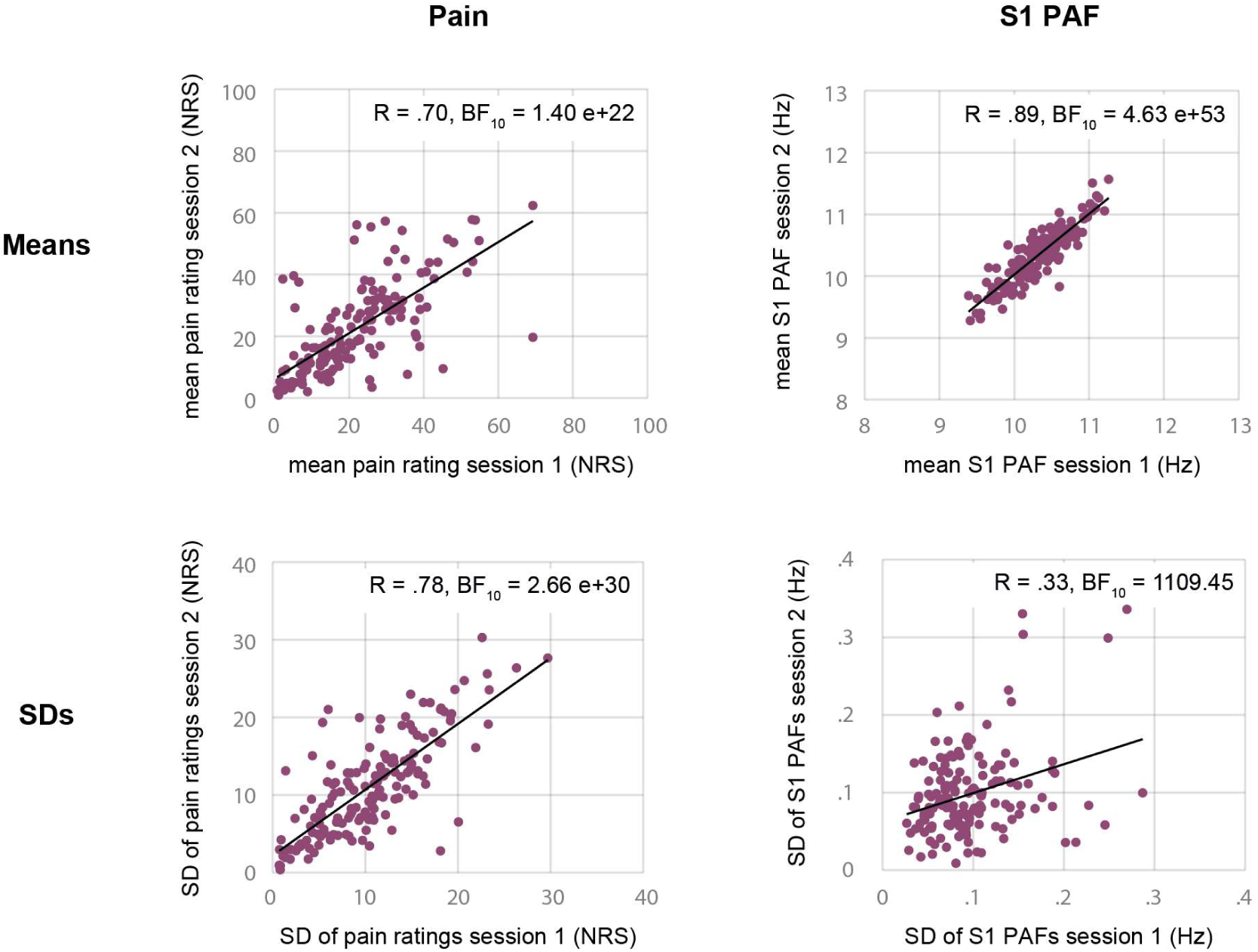
Between-session Bayesian linear correlations of single-participant means and SDs of pain ratings (left panels) and S1 resting-state PAFs during primary analyses (right panels). BF, Bayes factor; NRS, numerical rating scale; PAF, peak alpha frequency; R, Pearson correlation coefficient; S1, primary somatosensory cortex; SD, standard deviation.

**Supplementary Figure 2.**
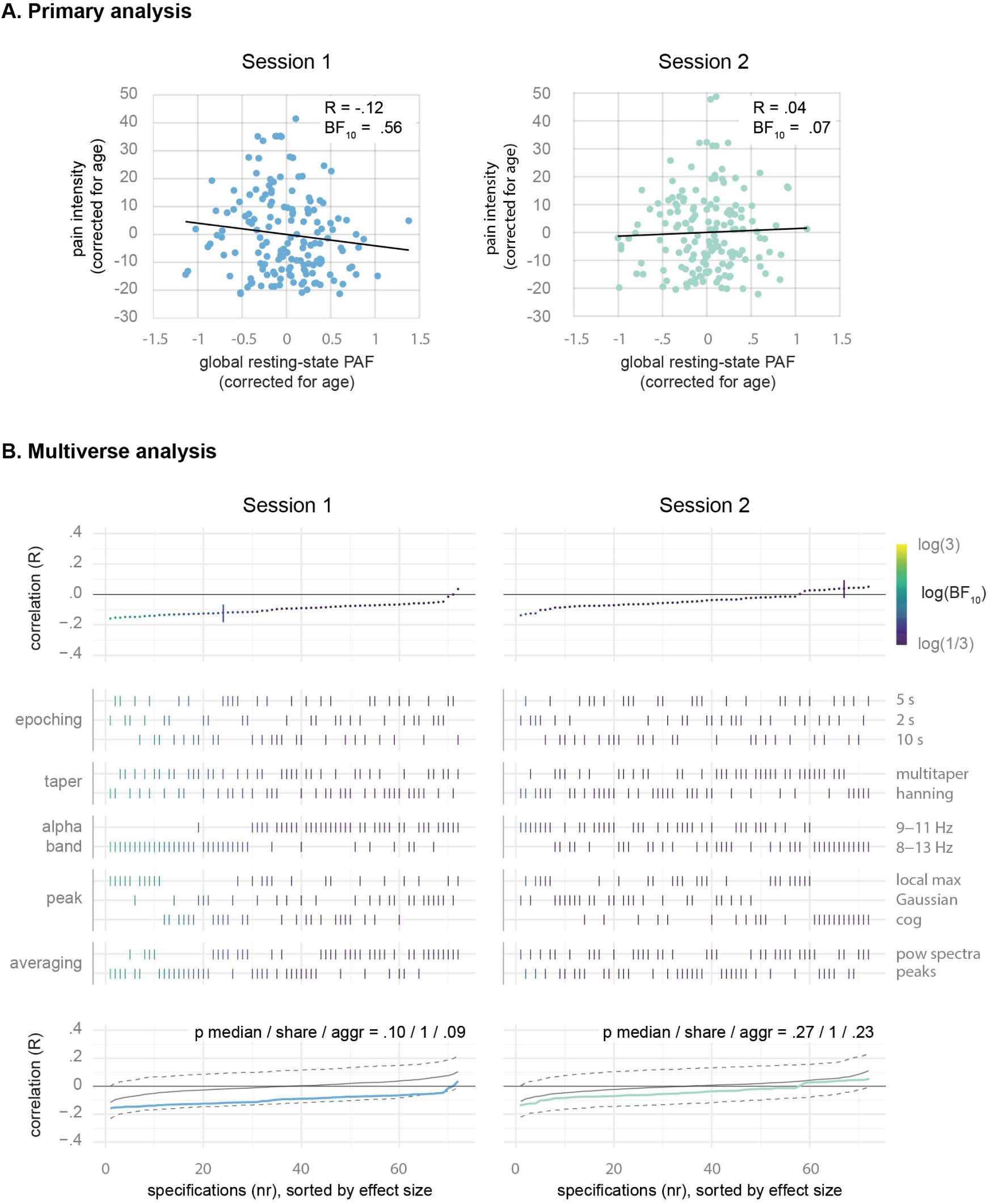
The relationship between global resting-state PAF and inter-individual variations of pain. A. Primary analysis. Scatter plots show the results of Bayesian linear correlations between global resting-state PAF and pain intensity, corrected for age. B. Multiverse analysis. The top panels show Pearson correlation coefficients obtained for all 72 specifications, sorted by effect size. The primary analysis is highlighted by a vertical bar. Middle panels mark the corresponding analytical choices of every specification (see Table 1). Results are color-coded based on Bayes factors. The scale’s upper and lower ends indicate moderate evidence for and against a relationship. The bottom panels display the inferential specification curve analysis showing the observed specification curve and the median and one-sided 95 % confidence intervals of 500 specification curves obtained from permutations of the data under the null hypothesis, all sorted by effect size. BF, Bayes factor; cog, center of gravity; PAF, peak alpha frequency; R, Pearson correlation coefficient.

**Supplementary Figure 3.**
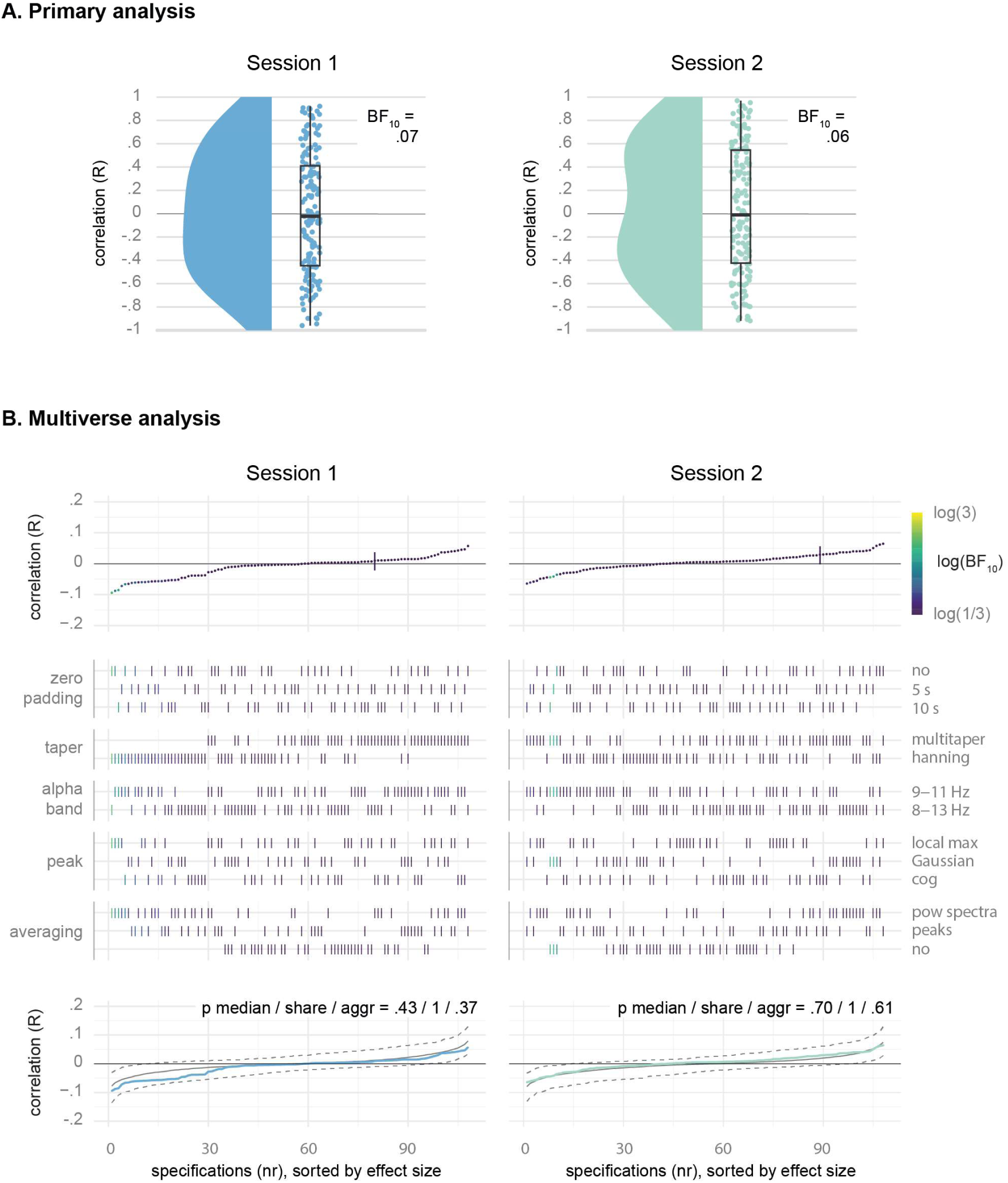
The relationship between global prestimulus PAF and intra-individual variations of pain. A. Primary analysis. Raincloud plots show single-participant correlation coefficients of the relation between global prestimulus PAF and pain intensity. Bayes factors are obtained from Bayesian t-tests of correlation coefficients vs. 0. B. Multiverse analysis. The top panels show Pearson correlation coefficients obtained for all 108 specifications, sorted by effect size. The primary analysis is highlighted by a vertical bar. Middle panels mark the corresponding analytical choices of every specification (see Table 1). Results are color-coded based on Bayes factors. The scale’s upper and lower ends indicate moderate evidence for and against a relationship. The bottom panels display the inferential specification curve analysis showing the observed specification curve and the median and one-sided 95 % confidence intervals of 500 specification curves obtained from permutations of the data under the null hypothesis, all sorted by effect size. BF, Bayes factor; cog, center of gravity; PAF, peak alpha frequency; R, Pearson correlation coefficient.

**Supplementary Figure 4.**
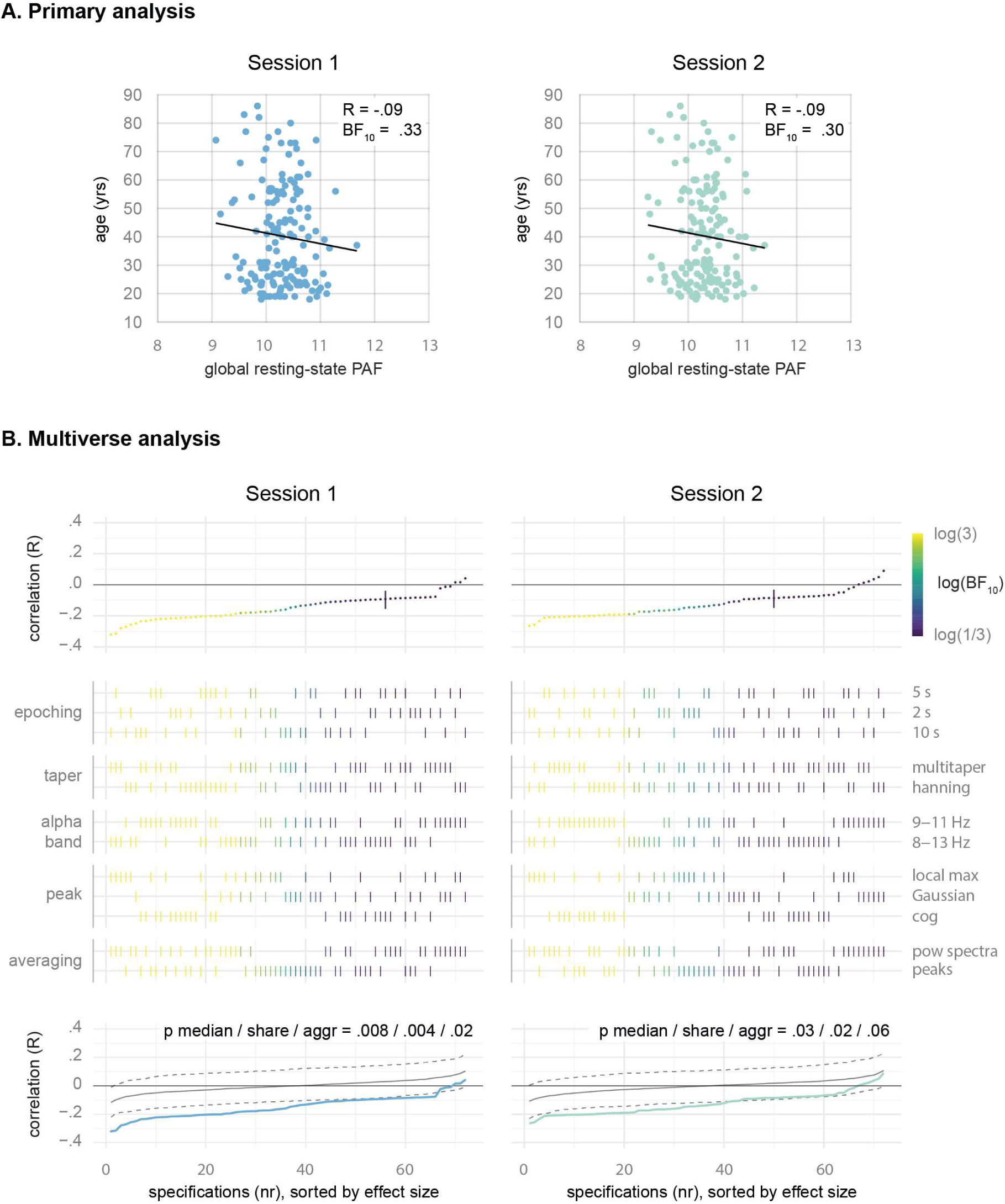
The relationship between global resting-state PAF and age. A. Primary analysis. Scatter plots show the results of Bayesian linear correlations between global resting-state PAF and age. B. Multiverse analysis. The top panels show Pearson correlation coefficients obtained for all 72 specifications, sorted by effect size. The primary analysis is highlighted by a vertical bar. Middle panels mark the corresponding analytical choices of every specification (see Table 1). Results are color-coded based on Bayes factors. The scale’s upper and lower ends indicate moderate evidence for or against a relationship. The bottom panels display the inferential specification curve analysis showing the observed specification curve and the median and one-sided 95 % confidence intervals of 500 specification curves obtained from permutations of the data under the null hypothesis, all sorted by effect size. BF, Bayes factor; cog, center of gravity; PAF, peak alpha frequency; R, Pearson correlation coefficient.

## Notes

### Competing Interest Statement

The authors have declared no competing interest.

